# Noninvasive thigh temperature mapping after cold water immersion and subsequent exercise using magnetic resonance spectrometry

**DOI:** 10.64898/2026.03.31.714134

**Authors:** Dorian Giraud, Arnaud Hays, Margaux Nussbaumer, Elodie Kopp, Nadège Corbin, Yann Le-Fur, Jean-Laurent Gardarein, Valéry Ozenne

## Abstract

Heat-related illnesses pose a significant public health challenge in Europe, resulting in increased mortality. Although cold water immersion (CWI) is the most effective treatment for heat stroke, its clinical use is limited. A better understanding of temperature changes in the peripheral body regions can lead to more effective CWI application. Nevertheless, most muscle temperature measurement techniques are invasive. This study evaluated magnetic resonance spectroscopy (MRS) for non-invasive assessment of intramuscular temperature during cold stress and rewarming.

Nine healthy volunteers (7 men, 2 women) participated in three 3T MRI sessions: baseline (PRE), immediately after 15 minutes of CWI at 10 degrees to the iliac crest (POST-CWI), and following 100-Watt cycling (POST-cycling). Each scan session included T1w and localized spectroscopy acquisitions in the right thigh. Absolute temperature was estimated from the proton resonance frequency shift between water and creatine peaks. The measurements were split into three groups of voxels, defined as follows: close to the top (TL), bottom (BL), or central (DL) thigh positions.

Measurement depth showed a location main effect (p<0.001, ηp^2=0.40), with DL (35.4[5.9] mm) significantly deeper than TL (22.5[4.2] mm) and BL (25.3[5.1] mm), remaining constant across phases. Temperature decreased significantly from PRE to POST-CWI across all locations (TL: p<0.001, d=2.74; BL: p<0.001, d=1.84; DL: p<0.005, d=1.14). Post-cycling temperature increased at all sites compared to POST-CWI (DL: p=0.040, d=1.06; TL: p<0.001, d=1.7; BL: p<0.001, d=1.80), though TL remained lower than PRE (p<0.017, d=1.48). During POST-CWI, DL showed a significantly higher temperature than TL (p<0.001, d=2.13) and BL (p<0.001, d=2.06).

These findings demonstrate that MRS-based temperature mapping provides unique anatomical and thermal characterization of muscle during thermoregulatory stress. While results are promising for understanding CWI mechanisms, validation in larger cohorts is necessary to establish clinical reliability and reproducibility for heat illness management.

## Introduction

We are currently living through a period of transition related to climate change. The IPCC’s AR6 report states that “many regions are expected to experience an increase in the likelihood of events associated with increased global warming, such as simultaneous heat waves and droughts, floods and wildfires” (Calvin et al., 2023). The increase in the number and intensity of heat waves will lead to a significant increase in deaths among vulnerable populations (Matthews et al., 2025). Preventing thermoregulatory disorders and introducing therapeutic solutions to cope with environmental stress could become a priority for health systems. Humans have a number of natural mechanisms that allow us to effectively fight the cold (Mäkinen, 2007), but maintaining thermoregulation in a hot, humid environment is more difficult. Indeed, the combination of the thermal hygrometry of the environment and the phenomena of convection and thermal radiation determine the human thermoregulatory capacity (Cramer & Jay, 2016). The two main mechanisms to combat heat are sweating and vasodilation, but when these mechanisms are overwhelmed, victims enter a stage of hyperthermia. The pathological stages range from mild “heat rash” to severe “heat stroke” (Barletta et al., 2024). This life-threatening emergency is classified as “classical” when it results from passive exposure to extreme ambient heat, or “exertional” in the context of intense physical exercise, and is currently associated with a mortality rate of 26.5% and 63.2% after admission to intensive care (Barletta et al., 2024). Cold water immersion is currently the most effective technique for treating heat stroke (Barletta et al., 2024). Treatment of heatstroke within the first 30 minutes has been shown to significantly reduce the risk of death, with the aim of achieving a cooling rate greater than 0.15°C per minute (Filep et al., 2020). Several numerical models have been developed to predict changes in body temperature during exposure to cold (Castellani et al., 2021; Gulati et al., 2022; Jacques et al., 2024). This could allow emergency services to accurately predict the dose of cold to be applied to the victim, depending on their physique, without causing pathologies associated with cold water immersion (Tipton et al., 2017). However, these models lack experimental validation (Xu et al., 2023), largely due to the challenge of measuring internal temperatures. Magnetic resonance imaging (MRI) overcomes these limitations by providing excellent soft tissue contrast for 3 dimension (3D) morphological analysis and functional imaging to quantify physiological changes. Techniques like T1w DIXON distinguish fat and water content, while MR angiography maps arterial and venous systems. MRI also enables functional assessments, such as blood perfusion via Intravoxel Incoherent Motion and fluid dynamics using 4D-flow. Unique among imaging modalities, MRI can non-invasively measure body temperature through localized or volumetric thermal mapping (Odéen & Parker, 2019).

The dependency of the proton resonance frequency shift on the temperature is the basis of the MR thermal monitoring (Odéen & Parker, 2019). It is used in two categories of techniques. First, relative measurement using phase mapping is the gold standard in interventional MRI during thermoablation therapies (De Senneville et al., 2007; Öcal et al., 2024). Phase mapping is designed to monitor large temperature changes (up to + 40 °C) during a limited period of time (∼5 min) to map the kinetics (1-8 s resolution) and spatial delivery (2 mm^3^ resolution) of energy. Second, proton MR spectrometry (MRS), which typically provides information on chemical composition or concentration, is the most common approach to extract absolute temperature using metabolites as an independent internal reference frequency (Odéen & Parker, 2019). The robustness of both methods is impaired by many perturbations such as motion, flow or magnetic field variations.

Two studies on thermoregulation were conducted using phase mapping to investigate heat deposition with radio frequence heating, specific absorption rate (SAR), heat dissipation in the foreleg (Simonis et al., 2016, 2017). However, as the phase mapping method is highly sensitive to fluctuations in the B0 field, it can only operate over a limited period of time and is only valid for measuring relative variations in temperature during the same MRI exams. At the same time, there has been renewed interest in MRS, as evidenced by recent clinical studies on changes in brain temperature. (Rzechorzek et al., 2022; Sharma et al., 2025). The major advantage is its ability to non-invasively measure the absolute temperature of tissues using either localized (Sung et al., 2022) or volumetric acquisitions (Thrippleton et al., 2014). While absolute MR-temperature measurements mostly target brain temperature changes and the impact of various neurological diseases such as epilepsy (Mueller et al., 2024; Sharma & Szaflarski, 2024), MRI is a promising technology for studying the impact of hypothermia, hyperthermia or climate change on the nervous system (Sisodiya et al., 2024; Taylor et al., 2016). Recently, Xiang Ren Tan et al, were able to show an increase in brain temperature under severe heat exposure (Tan et al., 2024).

The study aimed to evaluate the relevance of MRS for the non-invasive measurement of thigh intramuscular temperature in volunteers exposed to external thermal stress, including cold water immersion (CWI) and subsequent warm-up cycling exercise.

## Method

Two female and seven male volunteers (mean ± SD; age: 37 ± 13 years, body mass: 73 ± 12 kg; height: 175 ± 6 cm; IMC: 23.4 ± 2.8, thigh volume: 3074,5 ± 660,1 cm^3^, fat volume: 808,8 ± 315,5 cm^3^) were recruited for this study. All participants were fully informed of the potential risks and requirements of the study and provided written informed consent. The study with approval granted by the local ethics committee (MAPIRM, RCB 2016-A00434-47, n°4) and the informed written consent of the participants. Physical characteristics of the volunteers are summarized in supplementary Table 1.

### Experimental design

Events of a single test session with representative pictures of each step of the procedure are indicated in Figure 1. The protocol was divided into 3 distinct steps, each interspersed with MRI scans. On arrival at the laboratory, participants underwent an initial MRI scan (referred as PRE-CWI) to establish baseline muscle temperature, followed by cold water immersion (CWI) at 10°C for 15 minutes at the level of the iliac crest (CWI dose of 1.5). On completion of the immersion protocol, the participant’s legs were dabbed dry (so as not to stimulate blood flow) and quickly transferred to the MRI in a wheelchair and allowed a period of passive recovery during a second MRI scan (referred as POST-CWI, up to 40 min). This was followed by 10 minutes of cycling exercise at 100 Watt, followed by a second passive recovery period during the final MRI scan (referred as POST-CYCLING, up to 35 min). The water temperature was continuously monitored and kept at 10(0.5) °C using a water temperature controller (CryopackPerf, Cryo control, Castanet Tolosan, France). The water was not stirred manually, but the cooling system necessarily involved suction and discharge of water into the basin, resulting in a movement of water in the bath. The cycling part was realised with cycling ergometer (TACX FLUX S, Tacx, Netherland).

**Figure 1:**
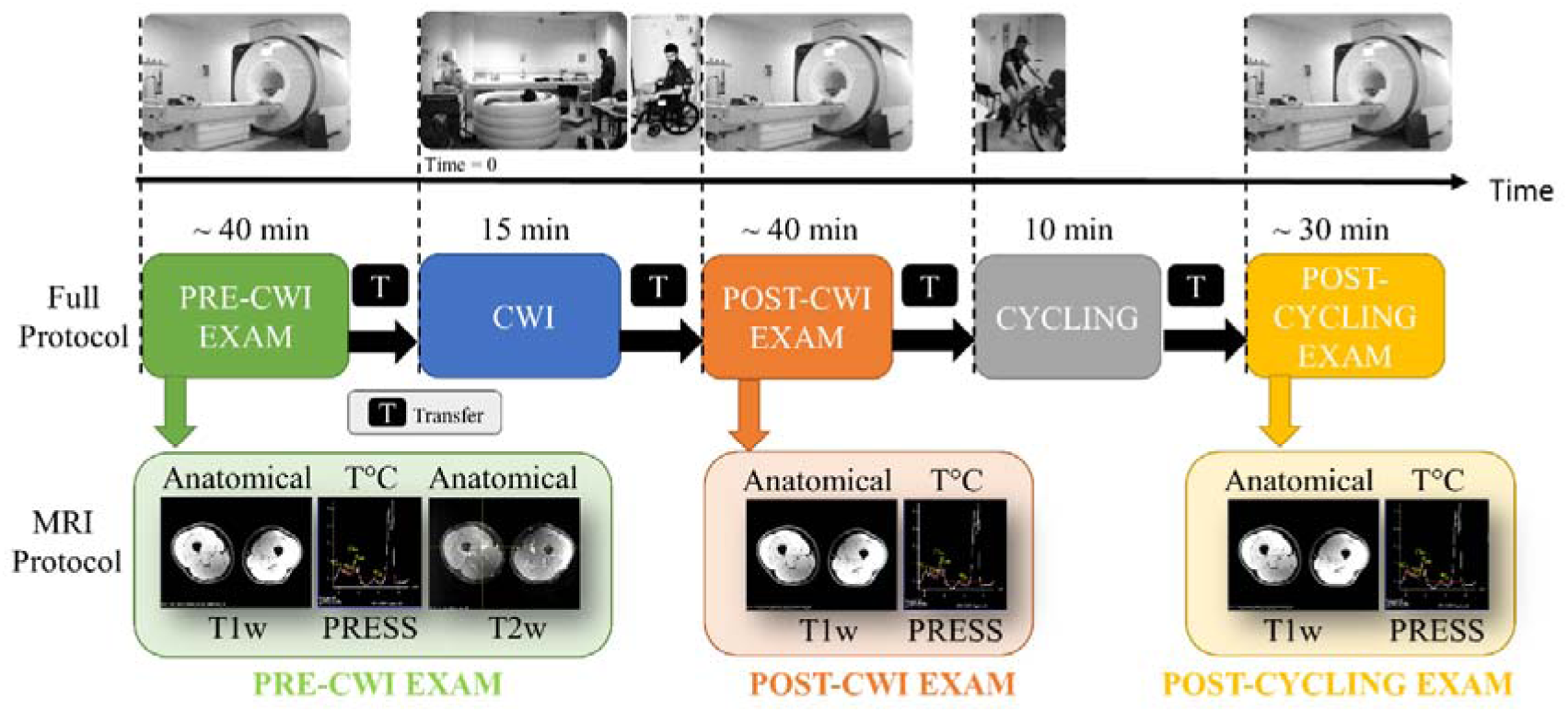
Experimental design. (Top) Events of a single session with representative pictures of each step of the procedure. (Middle) Schematic view and the approximate times of each MRI exam and of the transfer time. (Bottom) Simplified schematic view of the MRI acquisitions for each phase. The person in the photos is one of the authors.

#### MRI acquisitions

For each session, the volunteer lies on the table so that the right leg is at the isocenter of the magnet. MRI scans were recorded at 3 T (MAGNETOM Prisma, Siemens Healthineers, Erlangen, Germany) at the thigh levels using a spine coil on the bottom and one body matrix coil on the top of the thigh. To avoid thermal insulation of the thigh, the thigh was slightly raised with a cushion so that it was not in contact with the MRI table. However, the body matrix coil was in contact with the anterior part of the thigh as shown in Figure 2.

**Figure 2:**
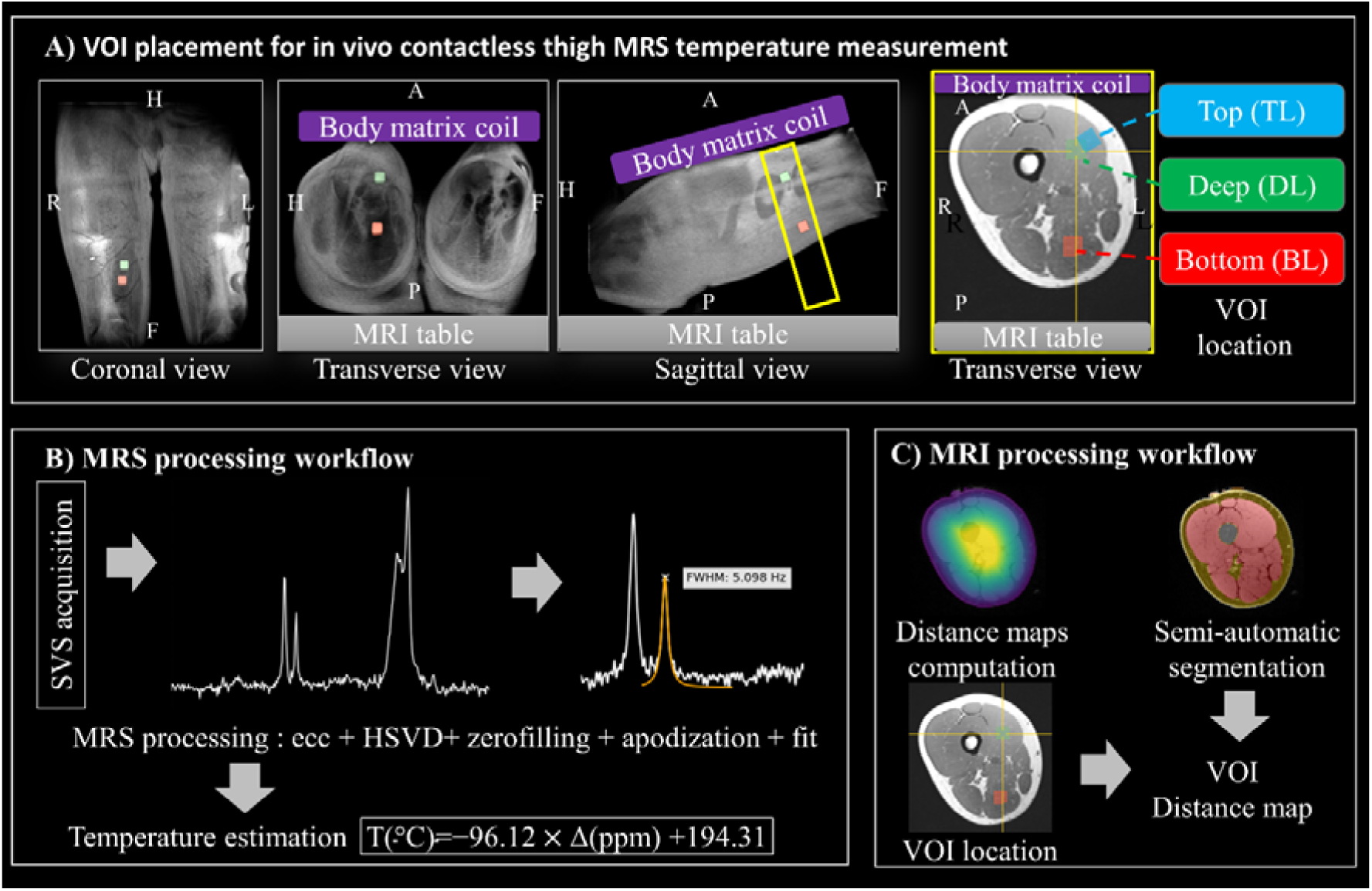
(A) Representative MRS VOI placement in the right thigh of volunteer #5 using 3D volume rendering of the T1-weighted Dixon in-phase image in coronal, transverse and sagittal views. To avoid thermal insulation of the thigh, the thigh was slightly raised so that it was not in contact with the MRI table (shown in green). However, the receiving antenna (shown in purple) was in contact with the anterior part of the thigh. Axial section of T1-weighted Dixon in-phase image with VOI location. The measurements were split into three groups of voxels defined such as: close to the muscle surface (top location in blue) or in a more central position (deep location in green) in the vastus medialis and close to the muscle surface (bottom location) in the semitendinosus or semimembranosus. A=Anterior, P=Posterior, R=Right, L=Left, H=Head, F=Foot. (B) The absolute temperature T in °C was estimated from the frequency shift Δ in ppm between the water peak and the Creatine peak of the PRESS spectrum. MRS processing workflow includes eddy courant correction, water suppression using HSVD, zerofilling, apodization and Lorentzian fit of the creatine and water peaks. (C) MRI post-processing steps include segmentation distance map computation.

The reference examination includes a localiser followed by a T1-weighted (T1w) Dixon and a T2-weighted (T2w) Dixon sequences. The parameters of the two-point Dixon 3D gradient-echo (GRE) sequence were: TR = 4.08 msec; TE1= 1.31 msec; TE2 = 2.54 msec; bandwidth = 820 Hz/pixel; flip angle = 9°; matrix size = 320 x 320 x 192; resolution =1.34 x 1.34 x 1.3 mm^3^; field of view = 430 x 430 x 250 mm^3^; partial fourier ⅞; average number = 6; acquisition time: 5:58 min. The parameters of the Dixon TSE were: TR= 6100 ms; TE=74 ms; pixel bandwidth = 473 Hz/pixel; acquisition matrix, 210×320×20, resolution = 1.25 x 1.25 x 5.0 mm^3^; slice gap, 2.5 mm; field of view =, 262 x 400 x 150 mm^3^; TSE factor, 10; flip angle 160°, acceleration =2; averages, 1; acquisition time: 2:10 min. Water-fat separation of the T1w and T2w images was performed online using the vendor’s algorithm.

The single voxel spectroscopy (SVS) technique was applied for the ^1^H MR spectroscopy temperature measurement method using a standard clinical Point-RESolved Spectroscopy (PRESS) sequence. Six preparation scans with water unsuppressed were first acquired followed by 128 averaged excitations. Parameters were: *TR* = 1200 ms, TE = 90 ms, bandwidth = 2000 Hz, flip angle = 65°, vector size =1024, volume of interest (VOI) of 12×12×12 mm^3^ to 10×10×10 mm^3^ (subject 8 and 9), acquisition time: 2:42 min. Six to eight localized spectroscopy sequences were then performed in locations defined in a position one-third of the way up the thigh (Figure 2). The measurements were thus split into three groups of voxels defined such as: close to the muscle surface (Top Location (TL) in blue) or in a more central position (Deep Location (DL) in green) in the vastus medialis and close to the muscle surface (Bottom Location (BL) in red) in the semitendinosus or semimembranosus (Figure 2).

The POST and CYCLE exam differed slightly from the reference examination. To limit the time between leaving the pool and the spectroscopy acquisitions, the T2w sequence was not applied and the number of averages in T1w Dixon sequence was reduced to 4 to have a shorter acquisition time equal to 1:51 min.

For the calibration procedure, a water bath system was used to heat a bottle containing water and the phantom tube from 28°C to 40°C. The temperature of the creatine gel and the water around it was monitored using two LumaSense optical fibers connected to a Luxtron812. Both of these temperatures were recorded and saved every second using a Raspberry Pi. A resting period was required before starting each 5-minute acquisition to achieve a satisfying temperature stability. Acquisitions were performed with the same SVS PRESS sequence using a 20-channel receive head coil on the same 3T scanner. The same parameters were used with the exception of: number of excitations = 256; acquisition time: 5:16 min.

#### Data processing

Muscle, fat and bone regions of the baseline exam were first segmented semi-automatically in 3DSlicer with the region growing tools using T1-weighted fat– and water-separated images. Then, the T1w images of the baseline exam were co-registered on the corresponding images of the post CWI and post exercise exam using ANTs [ref Avants2011] with antsRegistrationSyNQuick.sh and default parameters. Finally, label maps of tissue composition of the baseline exam were automatically mapped to the space of the T1w images of the post CWI and post exercise, preventing additional manual segmentation. Total, muscle and fat volume and ratio were computed using the corresponding label maps with ANTS using ‘LabelGeometryMeasures’. Distance maps of the thigh (with and without the peripheral fat) were computed using the Signed distance function of the SimpleITK library.

The absolute temperature T in °C was estimated from the distance (or frequency shift) Δshift in ppm between the water peak and the creatine peak. The Python library ‘Suspect’ (https://github.com/openmrslab/suspect) was used to post-process the DICOM files. Eddy current correction was applied on unsuppressed and water suppressed spectrum. As a consequence, all spectrums were re-referenced to 4.7 ppm. The Hankel Singular Value Decomposition (HSVD) method was used to remove the residual water peak. Then zerofilling (factor 4) and apodization (1.25) were applied. A Lorentz peak-fitting method was set up to fully automate maximum creatine pic detection and ensure reliability [ref Hartman 2016] and reproducibility. All data, code and associated scripts have been shared for reproducibility purposes (see Code and data availability section). Lastly, the ‘Suspect’ library was used to extract the voxel position from the localised spectroscopy and the average distance within the voxel from the skin or muscle tip.

The relationship between temperature and the chemical shift of water relative to creatine, established during the calibration procedure (see supplementary figure 1), is defined by the following linear equation:

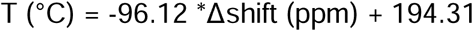

where T is the temperature (°C) and Δshift is the chemical shift difference between water and creatine (ppm). The model demonstrates high statistical significance (R = – 0.99, p < 0.001), with a root mean square error of 0.41°C and a mean absolute error of 0.33°C.

## Statistical analysis

On the temperature difference, a linear mixed model was performed using the ‘lmerTest’ (Kuznetsova et al., 2017) package in R due to its robustness in the result, as with the violation of the normality assumption (Schielzeth et al., 2020). To assess the location effect as a function of time, location and time were used as fixed effects and the random effects included only subject. Time corresponds to the 3-measurement time (Pre CWI, Post CWI and Post Exercise) described earlier in the method and location correspond to the 3 different voxels of measurement in the muscle. A pairwise comparison with Holm’s adjustment was then used for the interaction. The effect size (ES) of mean differences was determined using Cohen’s d coefficient, with effect sizes equal to or greater than 0.2, 0.6, and 1.2 interpreted as small, moderate, and large effects, respectively. Partial eta squared (ηp²) was calculated for each interaction on the result of lmer model.

## Results

All the result are presented as mean ± standard deviation (SD). The locations are called deep location (DL), top location (TL) and bottom location (BL). Temperature measurements were taken on average 1818 ± 639 sec (DL), 1805 ± 521 sec (TL) and 2377 ± 302 sec (BL) after the end of the CWI. The result for each location at each time is presented in Table 1

**Table 1:** The mean and standard deviation of the temperature and depth of measurement are shown for each location and measurement phase. SD: standard deviation; N: Number au measurement; BL: Bottom Location; DL: Deep Location; TL: Top Location

| Location | Time | Mean temperature (°C) | SD temperature (°C) | Mean depth (mm) | SD depth (mm) | N |
| --- | --- | --- | --- | --- | --- | --- |
| BL | PRE | 34.3 | 0.9 | 24.9 | 4.8 | 11 |
|  | POST | 31.7 | 1.7 | 25.4 | 5.4 | 14 |
|  | CYCLE | 34.2 | 0.8 | 25.6 | 5.3 | 12 |
| DL | PRE | 35.5 | 1.2 | 37.4 | 3.8 | 16 |
|  | POST | 34.3 | 0.9 | 32.3 | 6.2 | 21 |
|  | CYCLE | 35.2 | 0.7 | 37.0 | 5.8 | 20 |
| TL | PRE | 34.8 | 0.6 | 23.3 | 4.5 | 21 |
|  | POST | 31.7 | 1.5 | 21.3 | 4.1 | 19 |
|  | CYCLE | 33.9 | 0.6 | 22.2 | 3.7 | 13 |

### Depth differences

Depth showed only a location main effect (p<0.001, ηp²= 0.40). DL (35.4 ± 5.9 mm) was significantly higher than TL (22.5± 4.2 mm, p<0.001, d = 1.03) and than BL (25.3 ± 5.1 mm, p<0.001, d = 1.26). BL was higher than TL (p = 0.026, d =0.13). The location of each measurement zone did not change throughout the different phases of the protocol.

### Temperature effect

#### Time effect at each location

Figure 3 shows the effect of time at each location. The differences are shown using the compact letter display method for within-location comparisons and the star method for between-location comparisons. Temperature showed a significant Time x Location interaction (p<0.001; ηp²= 0.18). Temperature between PRE-CWI and POST-CWI decreased (TL: p < 0.001, d = 2.74; BL: p < 0.001, d = 1.84; DL: p < 0.005, d = 1.14). A significant increase of temperature was found after POST-CYCLING in comparison to POST-CWI (DL: p = 0.040, d = 1.06; TL: p < 0.001, d =1.7; BL: p < 0.001, d = 1.80)), regardless of the depth. POST CYCLING temperature was lower for TL in comparison to PRE-CWI (p < 0.017, d= 1.48).

**Figure 3:**
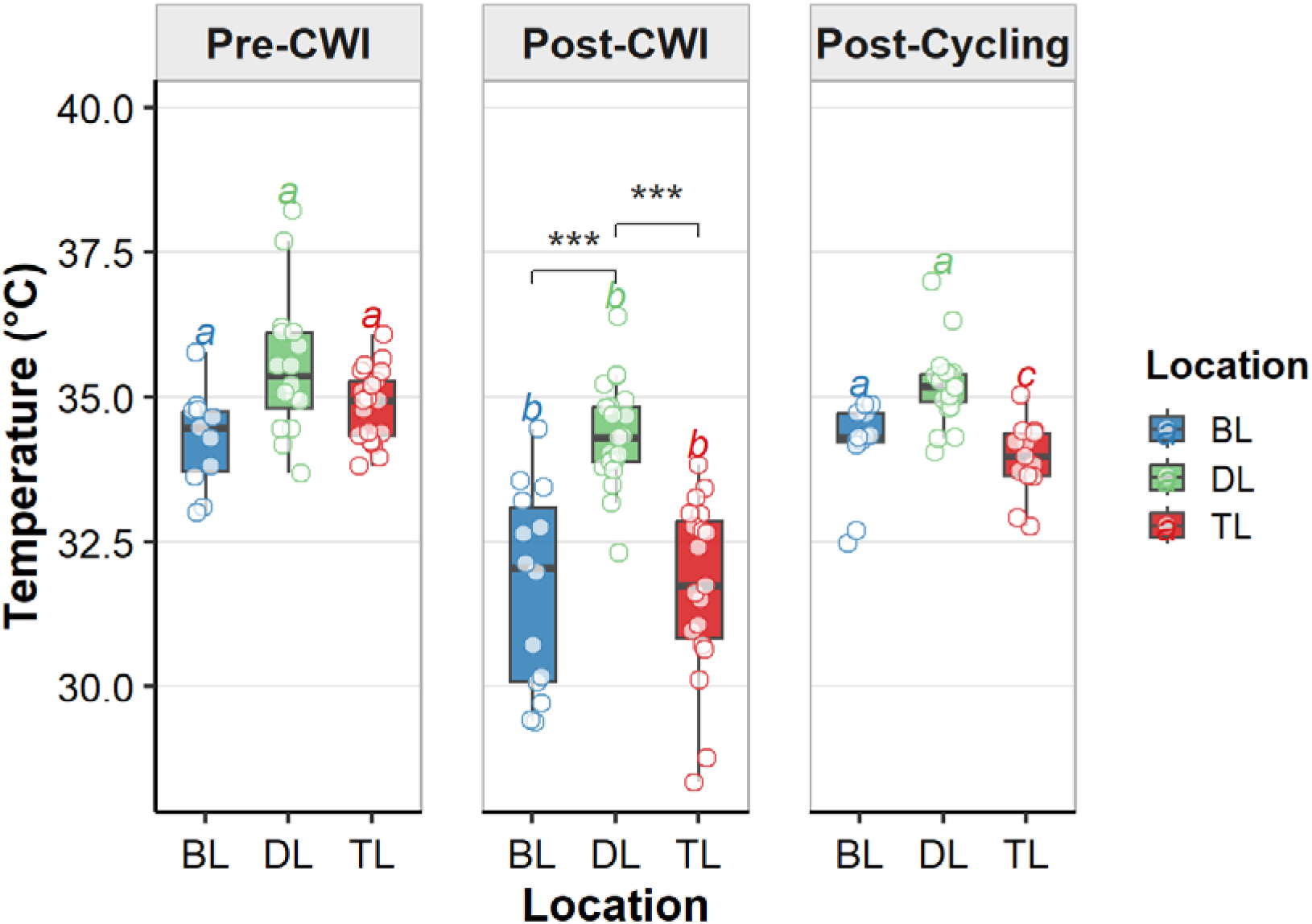
Temperature as a function of different exposure times. The differences between each location within each time are expressed as follows: ***→p ≤0.001. The differences between the different times within each location are expressed using the compact letter display statistical method. This allows only the significantly different values between them to be observed, without showing the p-value. Each data item is different if it does not contain at least one common letter. For example, A is significantly different from B, but not from AB. The p-value and the effect size of each difference are shown in the text. BL: Bottom Location; DL: Deep Location; TL: Top Location; CWI: Cold Water Immersion.

#### Location effect at each time

The effect of location at each time is shown in Figure 3 and the differences are shown using the star rating. Significant difference was found only during POST-CWI. DL was significantly higher than TL (p < 0.001, d = 2.13) and BL (p<0.001, d = 2.06).

### Example of single-subject results

Representative 1H-MRS spectrums across session of one subject are shown in Figure 4. For each case, a frequence shift could be observed between session with larger shift for the POST-CWI (orange curve). the spectroscopy-based estimated temperature for volunteer #3 and #5 are shown in Figure 5 and 6. Volunteer in Figure 5 showed less variability in temperature measurement than volunteer in Figure 6, particularly in BL position, despite equivalent depth across experimental phases.

**Figure 4:**
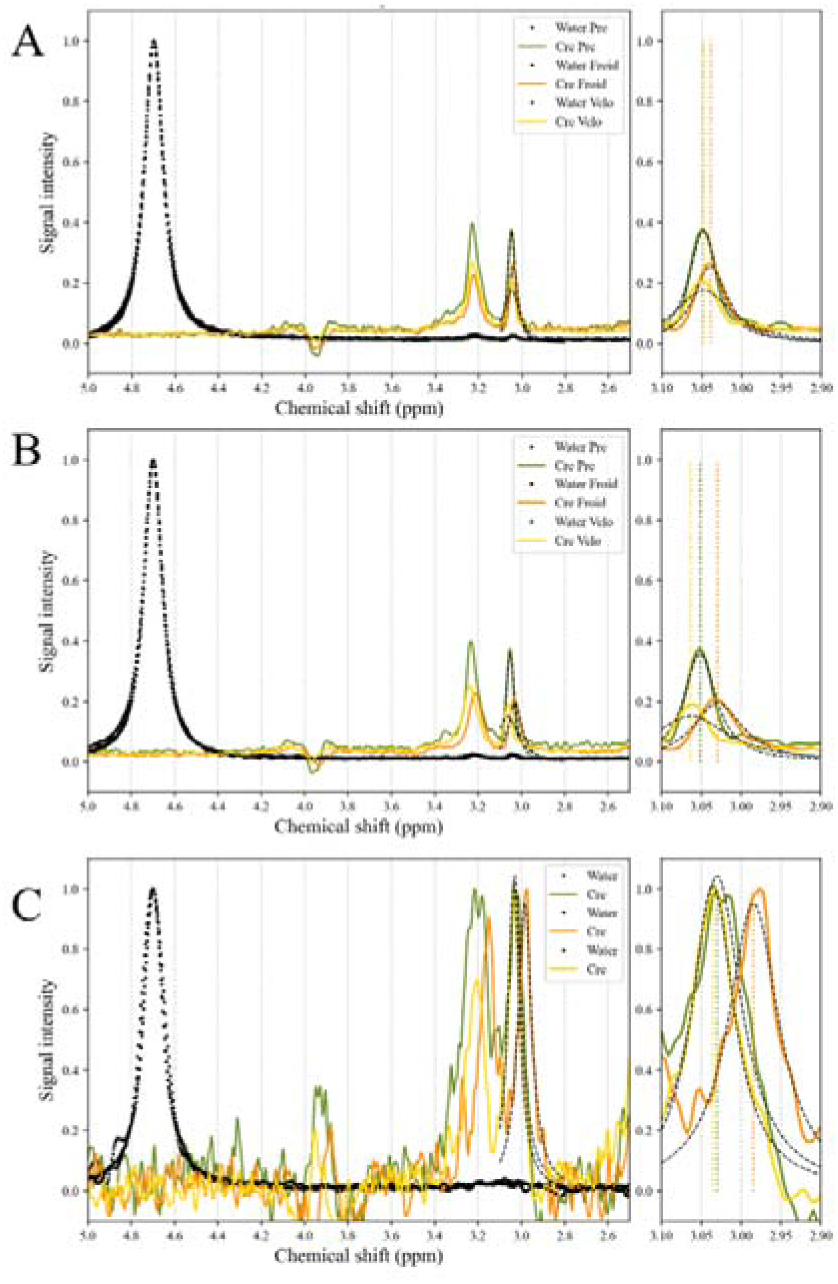
Representative 1H-MRS spectrum and frequency shift between session. Water unsuppressed spectrum are shown in black point and are all re-referenced to 4.7 ppm due to eddy current correction. As a consequence, chemical shift difference between water and creatine can be observed on the creatine peak. A Lorentz peak-fitting method was used to extract maximum creatine pic detection and applied in all session PRE-CWI (light green), POST-CWI (orange) and POST-CYCLING (yellow). A & B) Volonteer #3 C) Volonteer #5.

**Figure 5:**
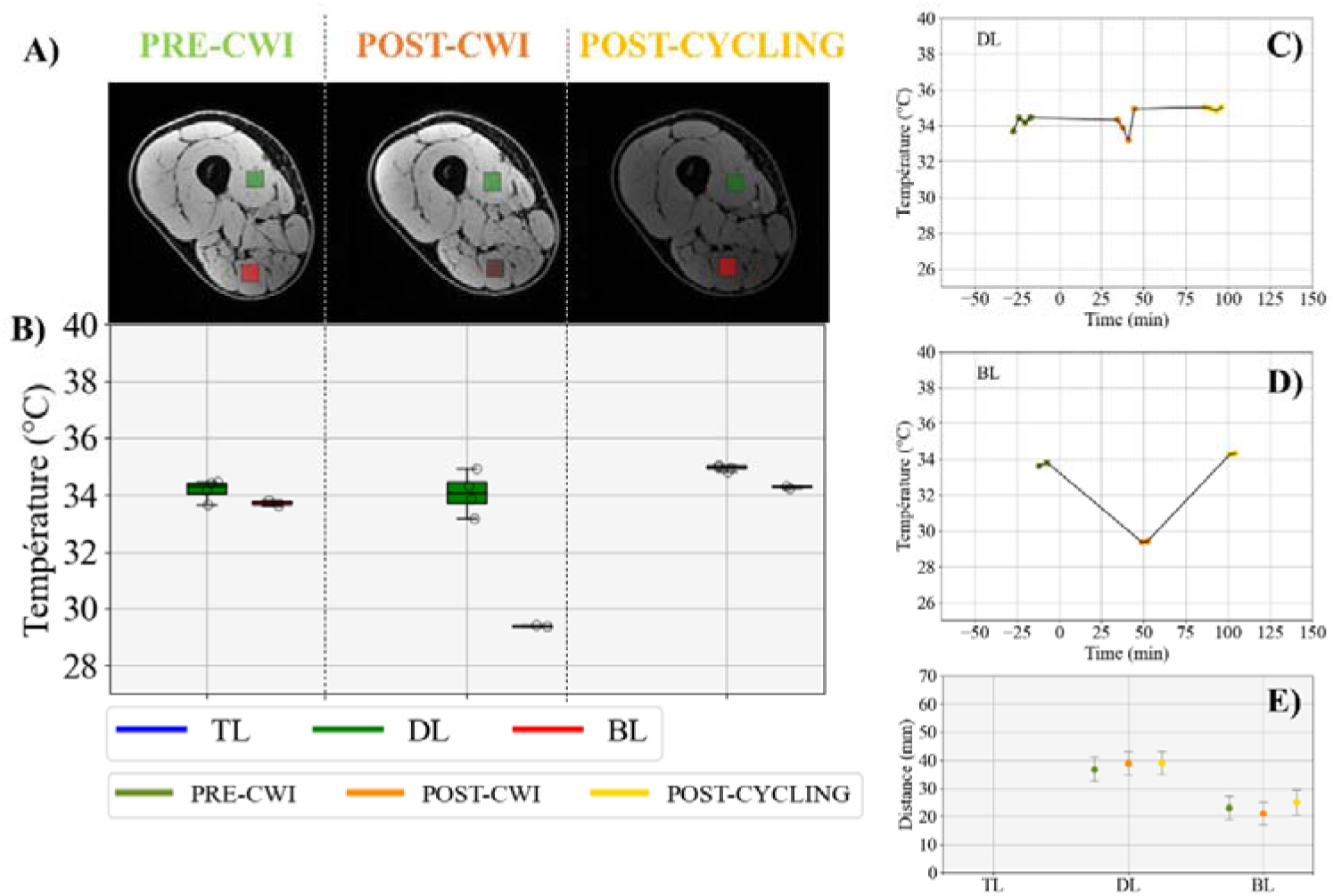
Contactless MRS-measured intramuscular temperature for volunteer #3. (A) VOI location for each phase placed in two locations overlaid on an axial T1 MRI image. One “center” VOI was placed in the vastus medialis (referred as DL, green) and the “lower” VOI was placed in the semimembranosus (referred as BL, red), no record was available in top location (TL). (B) Corresponding contactless MRS-measured intramuscular temperature PRE-CWI (light green), POST-CWI (orange) and POST-CYCLING (yellow). Data are presented as individual values and mean ± 95% CI for each group. (C, D) Contactless MRS-measured intramuscular temperature through time. Time was set at 0 at the beginning of CWI. (E) Mean ± Std of the distance maps computed from the skin surface in each VOI location and for each phase.

**Figure 6.**
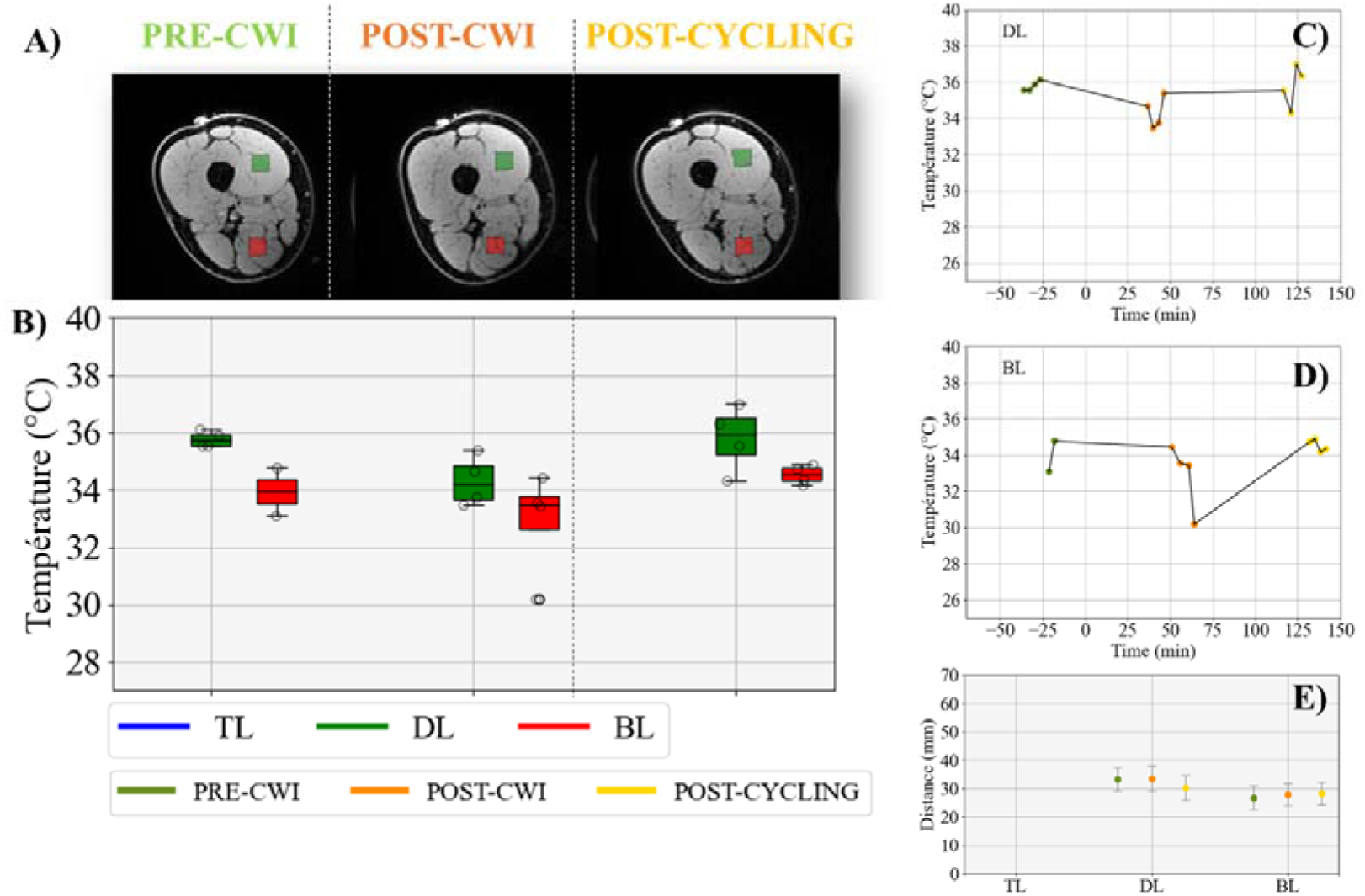
Contactless MRS-measured intramuscular temperature for volunteer #5. Same legends of figure 5 apply.

## Discussion

The aim of this study was to evaluate the relevance of MRS for non-invasive in vivo quantification of muscle temperature changes.

### MRI, a relevant modality for studying physiological responses to heat stress?

Compared to ultrasound, near-infrared spectroscopy or infrared thermography, MRI is a slow and complex modality for studying physiological responses to exercise. Volunteers must remain completely immobile in a supine position for between 30 minutes and 1 hour. Receiving channels are installed in the area to be imaged, which restricts access. Lastly, the presence of a strong magnetic field restricts the use of non-MRI compatible auxiliary equipment. Although cycling exercise within the bore has become common practice in MRI for muscle (Jeneson et al., 2010), cardiac (Pflugi et al., 2015) or brain assessment (Fontes et al., 2015), studies involving thigh thermal stress remain limited (Simonis et al., 2017).

One of our initial uncertainties regarding the protocol was the minimum time interval between the end of the CWI and the start of spectroscopic measurements. This period, which took around 30 minutes, included quick drying of the volunteer, transferring them to the scanner using a wheelchair to limit muscle activation, positioning the volunteer and the antennas and acquiring the anatomical images. Although, we were unable to measure the patient’s cooling kinetics, this timeframe was found compatible with the objectives as muscle temperature after CWI continued to fall until at least 1 hour after leaving the bath (Costello et al., 2012).

A second source of uncertainty was the positioning of the voxels by the MRI operator between each phases/session (Engelke et al., 2023). As the position of the voxels and their distance from the skin/air interface were likely to strongly influence the results, particular care was taken during the examination. Distance maps, and the mean ± SD distance values (see Table 1, Figures 3, 4 and figure supplementary 2) computed from the skin surface in each volume of interest (VOI) location and for each phase were, found acceptable with a SD close to 5 mm.

Spatial heterogeneity was observed among participants. Thigh volume can range from one to two times the original volume (x1.97 for volunteer #2 versus #7), while thigh fat volume can range from one to three times the original volume (x3.49 for volunteer #1 versus #3), as shown in Supplementary Figure 1. These variations can cause variability in the measured data.

Although single-voxel (1) H MRS using PRESS sequence is widely used for assessing absolute brain temperature in clinical studies (Cady et al., 1995; Childs et al., 2007; Rango et al., 2015; Whiteley et al., 2012, Thrippleton et al., 2014) the latter, consisting of 3 pulses (90, 180, 180), could induce a localised energy deposit. Here, a TR of 1200 ms was chosen to minimise this effect leading to a SAR of 0.1 W/kg. As a reminder, a SAR of 2 W/kg guarantees a maximum temperature increase of 0.5°C time-averaged over 6 minutes (according to the International Electrotechnical Commission: IEC 60601-2-33).

### Thigh temperature assessment

For each volunteer, localized spectroscopy sequences were repeated up to 4 times in each region and session. Temperature uncertainty was systematically found to be lower, in PRE-CWI and POST-CYCLING than in POST-CWI (see Table 1). Such result was visible either for each volunteer individually or for all volunteers together. For each volunteer, the uncertainty of the temperature during the POST-CWI session was bound to increase due to a stronger spatial gradient and a temporal evolution of the temperature. The kinetics involved being slower than the acquisition time (4× 2’30), temporal evolution of the temperature should have limited influence. While preprocessing and temperature quantification steps were fully scripted to ensure reproducibility as recommended (Near et al., 2021), considerable temperature variability was observed within most of the participants. Unphysical temperature changes were found as exemplified in figure 5, panel C with oscillation in PRE or abrupt temperature increase for the last point of POST or in figure 6, panel D with abrupt temperature increase for the first point of PRE. The limited spectral resolution could explain temperature oscillation, but other points are bound to the limit of the modality. Magnetic resonance spectroscopy has low sensitivity, which results in high measurement noise and is prone to artefact (Kreis, 2004).

Another hypothesis related to physiological responses could also be considered. Giraud et al. (2024) demonstrated that oxygen saturation in the vastus lateralis decreased approximately ten minutes after immersion in a cooling protocol identical to ours. As this decrease was not accompanied by a decrease in arterial blood flow, the authors concluded that it must be linked to an increase in energy metabolism associated with an increase in deoxygenated haemoglobin/myoglobin. However, the latter is a paramagnetic molecule (Pauling & Coryell, 1936), which increases the peak width at half height. This may have widened the creatine peak, leading to increased variability in the measurement points.

Mean temperature decrease after CWI was found up to 3.4°C, 2.9°C and 1.2°C in BL, TL and DL, so respectively around 25 mm, 23 mm and 35 mm, followed by a warming of the thigh after CYCLE for all depths. Even if BL measurement was taken on average 10 minutes later than DL and TL, points closer to the surface showed greater cooling than deeper points, as expected.

To our knowledge, only two studies have investigated muscle temperature evolution in five volunteers exposed to external thermal stress (Yoshioka et al., 2002, 2005). Measurements were taken during the cooling and warming phases using a device in contact with the skin. They were able to measure an absolute temperature drop of up to 26°C in the gastrocnemius muscle. However, the temperature measurements were subject to partial volume effect due to the voxel size of 20 to 40 ml. The estimated temperature values in the gastrocnemius and soleus muscles at ambient temperature were 33.6 ± 0.4 °C and 35.3 ± 0.4 °C, respectively. Nevertheless, the VOIs covered regions at different depths, limiting comparability with our study.

Two studies that looked at the temperature of the vastus lateralis muscle using thermocouples at rest found average temperatures of 34°C to 35°C at 2 cm depth and 35.7°C or 35.8°C at 3 cm depth (Costello et al., 2012; Mawhinney et al., 2020). In our study, we found equivalent temperatures at rest around these different depths (34.7 at 2 cm and 35.5 °C at 3 cm). However, several factors may influence the measurement. First, MR encodes information volumetrically by averaging the temperature across the voxel of interest and over the acquisition time. Therefore, the temperature measured by MRS may differ from the temperature measured by a punctual, instantaneous thermocouple in the presence of a high spatial gradient or temporal change. Secondly, as previously mentioned, the temperature derived from MRS is an indirect measurement that requires scanner-specific calibration (BabourinacBrooks et al., 2015; Sińczuk et al., 2025). Our calibration, as shown in supplementary figure 1 indicates an uncertainty of 0.4 °C. Our estimate assumes that the optical fibre has been perfectly calibrated.

As in the review by Vromans et al. (2019), a smaller decrease in muscle temperature was observed with increasing measurement depth after CWI. This can easily be explained by the phenomena of thermal diffusion within a solid body. Studies that employed the similar cooling protocol as ours found a temperature decrease of 1.4°C 9 minutes after immersion at a depth of 4 cm in the muscle (Broatch et al., 2014) or 5.2°C at a depth of 3 cm from the skin 1 minute after the end of CWI (Ihsan et al., 2014). The large difference in measurements between these two studies is probably due to the measurement distance, especially as Broatch et al. (2014) used the top layer of muscle as a reference point, whereas Ihsan et al. (2014) used the skin. Furthermore, in the study by Ihsan et al. (2014), only one leg was immersed, and the participants were in a standing position, unlike in our study and that of Broatch et al. (2014). The temperature decrease observed in the muscle at 3 cm appears to be lower than that measured by Ihsan et al. (2014). However, the latter immersed their leg after exercising, whereas we did not.

### Perspectives

Regarding the acquisition protocol, there is room for many improvements. 2D or 3D volumetric acquisition can be also performed using Magnetic Resonance Spectroscopic Imaging (MRSI) such as Echo-planar spectroscopic imaging (EPSI) (Rzechorzek et al., 2022; Weis et al., 2009) at the cost of a longer acquisition time (close to 15 minutes). These sequences would therefore be relevant for assessing resting temperature but may not be suitable in the presence of rapid temporal fluctuations. Recently, ultrafast magnetic resonance spectroscopic imaging producing high-resolution 3D 1 H metabolite maps and high-quality spatially resolved spectra (with a le nominal resolution of 2.4 × 2.4 × 3 mm3 in 5 minutes) have been reported (Lam et al., 2020) and may be of interest for in vivo volumetric 3D temperature mapping under thermal stress.

## Competing interests

The author(s) declare no competing interests.

## Code and data availability

Link to the calibration data is https://recherche.data.gouv.fr/fr (soon). Associated command scripts that compute in absolute temperature and command-line instructions for generating figures are available at this link https://github.com/Thermomics/CALMARS/. Link to the clinical data is https://recherche.data.gouv.fr/fr (soon). Associated command scripts that compute in absolute temperature, and that compute statistics (using a R script) and command-line instructions for generating figures are available at this link https://github.com/Thermomics/J_Peux_Pas_J_ai_Piscine.

## Author contributions

E.K., M.N, V.O. optimize the sequences, carried out MR experiments, collected the calibration data, and mrs analysis. D.G, E.K, J.L.G., V.O, conducted the mri-mrs study, developed the protocol, and carried out MR experiments. E.K, V.O, Y. L. F, N.C processed the mri-mrs data and D.G perform statistical analysis. D.G and V.O. drafted the manuscript. A.H, J.L.G. and V.O. jointly and equally participated in its design and coordination. All authors read and approved the final manuscript.

## Acknowledgements

The authors thank the volunteers who participated in this study as well as staff who assisted with data collection: Serge Anandra, Jérome Naulin and Dounia El Hamrani. The authors thank Equipex + HIPE ANR-21-ESRE-004.

## Notes

https://github.com/Thermomics/CALMARS/

https://github.com/Thermomics/J_Peux_Pas_J_ai_Piscine

### Competing Interest Statement

The authors have declared no competing interest.

## Bibliographie

1. Babourina-Brooks, B., Simpson, R., Arvanitis, T. N., Machin, G., Peet, A. C., & Davies, N. P. (2015). MRS thermometry calibration at 3 T: Effects of protein, ionic concentration and magnetic field strength. NMR in Biomedicine, 28(7), 792–800. 10.1002/nbm.3303

2. Barletta, J. F., Palmieri, T. L., Toomey, S. A., Harrod, C. G., Murthy, S., & Bailey, H. (2024). Management of Heat-Related Illness and Injury in the ICU: A Concise Definitive Review. Critical Care Medicine, 52(3), 362–375. 10.1097/CCM.0000000000006170

3. Broatch, J. R., Petersen, A., & Bishop, D. J. (2014). Postexercise Cold Water Immersion Benefits Are Not Greater than the Placebo Effect. Medicine & Science in Sports & Exercise, 46(11), 2139–2147. 10.1249/MSS.0000000000000348

4. Cady, E. B., D’Souza, P. C., Penrice, J., & Lorek, A. (1995). The Estimation of Local Brain Temperature by *in Vivo*^1^ H Magnetic Resonance Spectroscopy. Magnetic Resonance in Medicine, 33(6), 862–867. 10.1002/mrm.1910330620

5. Calvin, K., Dasgupta, D., Krinner, G., Mukherji, A., Thorne, P. W., Trisos, C., Romero, J., Aldunce, P., Barrett, K., Blanco, G., Cheung, W. W. L., Connors, S., Denton, F., Diongue-Niang, A., Dodman, D., Garschagen, M., Geden, O., Hayward, B., Jones, C., … Péan, C. (with Lee, H.). (2023). IPCC, 2023: Climate Change 2023: Synthesis Report. Contribution of Working Groups I, II and III to the Sixth Assessment Report of the Intergovernmental Panel on Climate Change [Core Writing Team, H. Lee and J. Romero (eds.)]. IPCC, Geneva, Switzerland. (First). Intergovernmental Panel on Climate Change (IPCC). 10.59327/IPCC/AR6-9789291691647

6. Castellani, M. P., Rioux, T. P., Castellani, J. W., Potter, A. W., & Xu, X. (2021). A geometrically accurate 3 dimensional model of human thermoregulation for transient cold and hot environments. Computers in Biology and Medicine, 138, 104892. 10.1016/j.compbiomed.2021.104892

7. Childs, C., Hiltunen, Y., Vidyasagar, R., & Kauppinen, R. A. (2007). Determination of regional brain temperature using proton magnetic resonance spectroscopy to assess brain–body temperature differences in healthy human subjects. Magnetic Resonance in Medicine, 57(1), 59–66. 10.1002/mrm.21100

8. Costello, J. T., Culligan, K., Selfe, J., & Donnelly, A. E. (2012). Muscle, Skin and Core Temperature after −110°C Cold Air and 8°C Water Treatment. PLoS ONE, 7(11), e48190. 10.1371/journal.pone.0048190

9. Cramer, M. N., & Jay, O. (2016). Biophysical aspects of human thermoregulation during heat stress. Autonomic Neuroscience, 196, 3–13. 10.1016/j.autneu.2016.03.001

10. De Senneville, B. D., Mougenot, C., Quesson, B., Dragonu, I., Grenier, N., & Moonen, C. T. W. (2007). MR thermometry for monitoring tumor ablation. European Radiology, 17(9), 2401–2410. 10.1007/s00330-007-0646-6

11. Engelke, K., Chaudry, O., Gast, L., Eldib, M. Ab., Wang, L., Laredo, J.-D., Schett, G., & Nagel, A. M. (2023). Magnetic resonance imaging techniques for the quantitative analysis of skeletal muscle: State of the art. Journal of Orthopaedic Translation, 42, 57–72. 10.1016/j.jot.2023.07.005

12. Filep, E. M., Murata, Y., Endres, B. D., Kim, G., Stearns, R. L., & Casa, D. J. (2020). Exertional Heat Stroke, Modality Cooling Rate, and Survival Outcomes: A Systematic Review. Medicina, 56(11), 589. 10.3390/medicina56110589

13. Fontes, E. B., Okano, A. H., De Guio, F., Schabort, E. J., Min, L. L., Basset, F. A., Stein, D. J., & Noakes, T. D. (2015). Brain activity and perceived exertion during cycling exercise: An fMRI study. British Journal of Sports Medicine, 49(8), 556–560. 10.1136/bjsports-2012-091924

14. Gulati, T., Hatwar, R., Unnikrishnan, G., Rubio, J. E., & Reifman, J. (2022). A 3-D virtual human model for simulating heat and cold stress. Journal of Applied Physiology (Bethesda, Md.: 1985), 133(2), 288–310. 10.1152/japplphysiol.00089.2022

15. Ihsan, M., Watson, G., Choo, H. C., Lewandowski, P., Papazzo, A., Cameron-Smith, D., & Abbiss, C. R. (2014). Postexercise Muscle Cooling Enhances Gene Expression of PGC-1α. Medicine & Science in Sports & Exercise, 46(10), 1900–1907. 10.1249/MSS.0000000000000308

16. Jacques, J., Rioux, T., Xu, X., & Castellani, J. (2024). Human Thermoregulation and Spatial Temperature for Frostbite Prediction with Bio-Heat Transfer Model. COMSOL conference.

17. Jeneson, J. A. L., Schmitz, J. P. J., Hilbers, P. A. J., & Nicolay, K. (2010). An MR-compatible bicycle ergometer for in-magnet whole-body human exercise testing. Magnetic Resonance in Medicine, 63(1), 257–261. 10.1002/mrm.22179

18. Kreis, R. (2004). Issues of spectral quality in clinical^1^ H magnetic resonance spectroscopy and a gallery of artifacts. NMR in Biomedicine, 17(6), 361–381. 10.1002/nbm.891

19. Kuznetsova, A., Brockhoff, P. B., & Christensen, R. H. (2017). lmerTest package: Tests in linear mixed effects models. Journal of Statistical Software, 82, 1–26.

20. Lam, F., Li, Y., Guo, R., Clifford, B., & Liang, Z. (2020). Ultrafast magnetic resonance spectroscopic imaging using SPICE with learned subspaces. Magnetic Resonance in Medicine, 83(2), 377–390. 10.1002/mrm.27980

21. Mäkinen, T. M. (2007). Human cold exposure, adaptation, and performance in high latitude environments. American Journal of Human Biology, 19(2), 155–164. 10.1002/ajhb.20627

22. Matthews, T., Raymond, C., Foster, J., Baldwin, J. W., Ivanovich, C., Kong, Q., Kinney, P., & Horton, R. M. (2025). Mortality impacts of the most extreme heat events. Nature Reviews Earth & Environment, 6(3), 193–210. 10.1038/s43017-024-00635-w

23. Mawhinney, C., Heinonen, I., Low, D. A., Han, C., Jones, H., Kalliokoski, K. K., Kirjavainen, A., Kemppainen, J., Di Salvo, V., Weston, M., Cable, T., & Gregson, W. (2020). Changes in quadriceps femoris muscle perfusion following different degrees of cold-water immersion. Journal of Applied Physiology, 128(5), 1392–1401. 10.1152/japplphysiol.00833.2019

24. Mueller, C., Hong, H., Sharma, A. A., Qin, H., Benveniste, E. N., & Szaflarski, J. P. (2024). Brain temperature, brain metabolites, and immune system phenotypes in temporal lobe epilepsy. Epilepsia Open, 9(6), 2454–2466. 10.1002/epi4.13082

25. Near, J., Harris, A. D., Juchem, C., Kreis, R., Marjańska, M., Öz, G., Slotboom, J., Wilson, M., & Gasparovic, C. (2021). Preprocessing, analysis and quantification in single-voxel magnetic resonance spectroscopy: Experts’ consensus recommendations. NMR in Biomedicine, 34(5), e4257. 10.1002/nbm.4257

26. Öcal, O., Dietrich, O., Lentini, S., Bour, P., Faller, T., Ozenne, V., Maier, F., Fabritius, M. P., Puhr-Westerheide, D., Schmidt, V. F., Öcal, E., Seidensticker, R., Wildgruber, M., Ricke, J., & Seidensticker, M. (2024). Predicting liver ablation volumes with real-time MRI thermometry. JHEP Reports, 6(11), 101199. 10.1016/j.jhepr.2024.101199

27. Odéen, H., & Parker, D. L. (2019). Magnetic resonance thermometry and its biological applications – Physical principles and practical considerations. Progress in Nuclear Magnetic Resonance Spectroscopy, 110, 34–61. 10.1016/j.pnmrs.2019.01.003

28. Pauling, L., & Coryell, C. D. (1936). The Magnetic Properties and Structure of Hemoglobin, Oxyhemoglobin and Carbonmonoxyhemoglobin. Proceedings of the National Academy of Sciences, 22(4), 210–216. 10.1073/pnas.22.4.210

29. Pflugi, S., Roujol, S., Akçakaya, M., Kawaji, K., Foppa, M., Heydari, B., Goddu, B., Kissinger, K., Berg, S., Manning, W. J., Kozerke, S., & Nezafat, R. (2015). Accelerated cardiac MR stress perfusion with radial sampling after physical exercise with an MR-compatible supine bicycle ergometer. Magnetic Resonance in Medicine, 74(2), 384–395. 10.1002/mrm.25405

30. Rango, M., Bonifati, C., & Bresolin, N. (2015). Post-Activation Brain Warming: A 1-H MRS Thermometry Study. PLOS ONE, 10(5), e0127314. 10.1371/journal.pone.0127314

31. Rzechorzek, N. M., Thrippleton, M. J., Chappell, F. M., Mair, G., Ercole, A., Cabeleira, M., The CENTER-TBI High Resolution ICU (HR ICU) Sub-Study Participants and Investigators, Rhodes, J., Marshall, I., & O’Neill, J. S. (2022). A daily temperature rhythm in the human brain predicts survival after brain injury. Brain, 145(6), 2031–2048. 10.1093/brain/awab466

32. Schielzeth, H., Dingemanse, N. J., Nakagawa, S., Westneat, D. F., Allegue, H., Teplitsky, C., Réale, D., Dochtermann, N. A., Garamszegi, L. Z., & Araya-Ajoy, Y. G. (2020). Robustness of linear mixed-effects models to violations of distributional assumptions. Methods in Ecology and Evolution, 11(9), 1141–1152. 10.1111/2041-210X.13434

33. Sharma, A. A., Nenert, R., & Szaflarski, J. P. (2025). Finding the Fire: Mapping Brain Temperature Elevations as a Surrogate of Focal Neuroinflammation. In D. Sone (Ed.), Molecular Imaging for Brain Diseases (Vol. 222, pp. 127–147). Springer US. 10.1007/978-1-0716-4494-2_9

34. Sharma, A. A., & Szaflarski, J. P. (2024). The longitudinal effects of cannabidiol on brain temperature in patients with treatment-resistant epilepsy. Epilepsy & Behavior, 151, 109606. 10.1016/j.yebeh.2023.109606

35. Simonis, F. F. J., Petersen, E. T., Lagendijk, J. J. W., & Van Den Berg, C. A. T. (2016). Feasibility of measuring thermoregulation during RF heating of the human calf muscle using MR based methods. Magnetic Resonance in Medicine, 75(4), 1743–1751. 10.1002/mrm.25710

36. Simonis, F. F. J., Raaijmakers, A. J. E., Lagendijk, J. J. W., & Van Den Berg, C. A. T. (2017). Validating subject-specific RF and thermal simulations in the calf muscle using MR-based temperature measurements: Validating RF and Thermal Simulations with MRI Measurements. Magnetic Resonance in Medicine, 77(4), 1691–1700. 10.1002/mrm.26244

37. Sińczuk, M., Rogala, J., & Bogorodzki, P. (2025). MRS thermometry – Importance of scanner-specific calibrations for accurate brain temperature estimations. Biocybernetics and Biomedical Engineering, 45(3), 451–456. 10.1016/j.bbe.2025.06.001

38. Sisodiya, S. M., Gulcebi, M. I., Fortunato, F., Mills, J. D., Haynes, E., Bramon, E., Chadwick, P., Ciccarelli, O., David, A. S., De Meyer, K., Fox, N. C., Davan Wetton, J., Koltzenburg, M., Kullmann, D. M., Kurian, M. A., Manji, H., Maslin, M. A., Matharu, M., Montgomery, H., … Hanna, M. G. (2024). Climate change and disorders of the nervous system. The Lancet Neurology, 23(6), 636–648. 10.1016/S1474-4422(24)00087-5

39. Tan, X. R., Stephenson, M. C., Alhadad, S. B., Loh, K. W. Z., Soong, T. W., Lee, J. K. W., & Low, I. C. C. (2024). Elevated brain temperature under severe heat exposure impairs cortical motor activity and executive function. Journal of Sport and Health Science, 13(2), 233–244. 10.1016/j.jshs.2023.09.001

40. Taylor, L., Watkins, S. L., Marshall, H., Dascombe, B. J., & Foster, J. (2016). The Impact of Different Environmental Conditions on Cognitive Function: A Focused Review. Frontiers in Physiology, 6. 10.3389/fphys.2015.00372

41. Thrippleton, M. J., Parikh, J., Harris, B. A., Hammer, S. J., Semple, S. I. K., Andrews, P. J. D., Wardlaw, J. M., & Marshall, I. (2014). Reliability of MRSI brain temperature mapping at 1.5 and 3 T. NMR in Biomedicine, 27(2), 183–190. 10.1002/nbm.3050

42. Tipton, M. J., Collier, N., Massey, H., Corbett, J., & Harper, M. (2017). Cold water immersion: Kill or cure?: Cold water immersion: kill or cure? Experimental Physiology, 102(11), 1335–1355. 10.1113/EP086283

43. Vromans, B. A., Thorpe, R. T., Viroux, P. J., & Tiemessen, I. J. (2019). Cold water immersion settings for reducing muscle tissue temperature: A linear dose-response relationship. The Journal of Sports Medicine and Physical Fitness, 59(11). 10.23736/S0022-4707.19.09398-8

44. Weis, J., Covaciu, L., Rubertsson, S., Allers, M., Lunderquist, A., & Ahlström, H. (2009). Noninvasive monitoring of brain temperature during mild hypothermia. Magnetic Resonance Imaging, 27(7), 923–932. 10.1016/j.mri.2009.01.011

45. Whiteley, W. N., Thomas, R., Lowe, G., Rumley, A., Karaszewski, B., Armitage, P., Marshall, I., Lymer, K., Dennis, M., & Wardlaw, J. (2012). Do acute phase markers explain body temperature and brain temperature after ischemic stroke? Neurology, 79(2), 152–158. 10.1212/WNL.0b013e31825f04d8

46. Xu, X., Rioux, T. P., & Castellani, M. P. (2023). Three dimensional models of human thermoregulation: A review. Journal of Thermal Biology, 112, 103491. 10.1016/j.jtherbio.2023.103491

47. Yoshioka, Y., Oikawa, H., Ehara, S., Inoue, T., Ogawa, A., Kambara, Y., Itazawa, S.-I., & Kubokawa, M. (2002). Noninvasive Estimation of Temperature and pH in Human Lower Leg Muscles using^1^ H Nuclear Magnetic Resonance Spectroscopy. Journal of Spectroscopy, 16(3–4), 183–190. 10.1155/2002/392987

48. Yoshioka, Y., Oikawa, H., Ehara, S., Inoue, T., Ogawa, A., Kanbara, Y., & Kubokawa, M. (2005). Noninvasive measurement of temperature and fractional dissociation of imidazole in human lower leg muscles using^1^ H-nuclear magnetic resonance spectroscopy. Journal of Applied Physiology, 98(1), 282–287. 10.1152/japplphysiol.00437.2004

