## Supplementary_figures for "Noninvasive thigh temperature mapping after cold water immersion and subsequent exercise using magnetic resonance spectrometry"

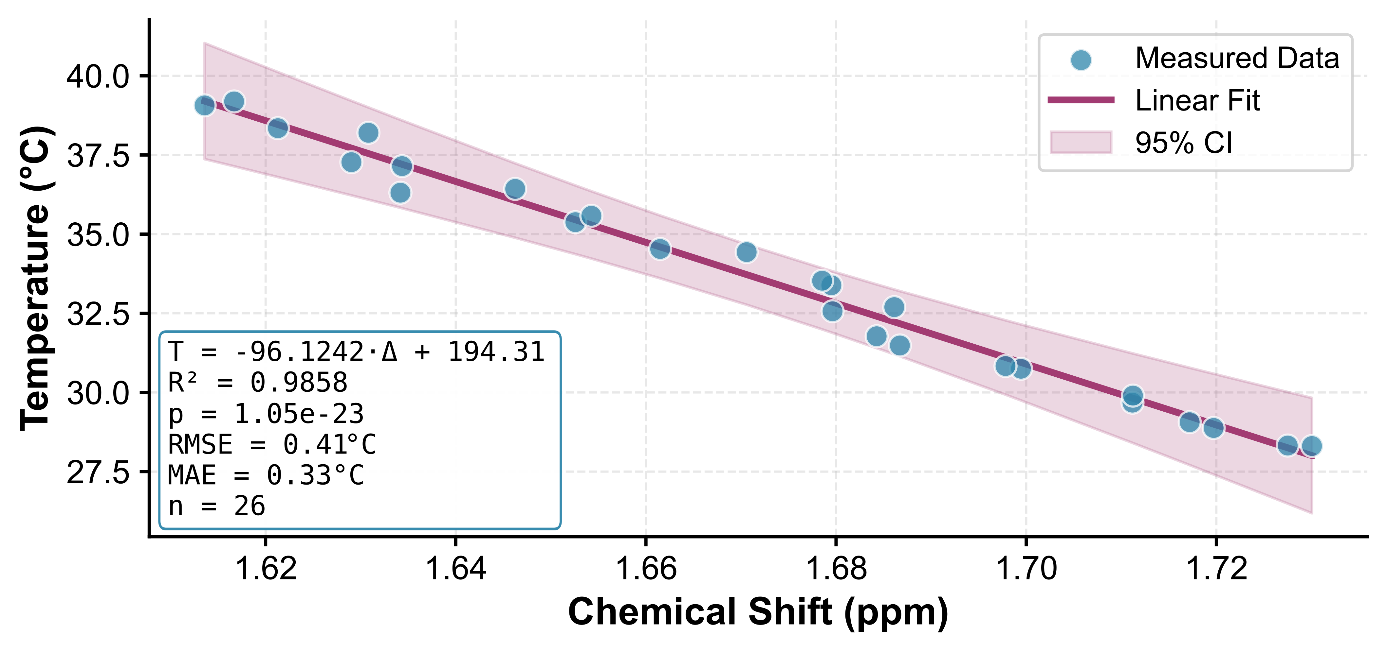

Supplementary figure 1. Temperature from optical fibers is plotted as function of chemical shift for connected. Temperature calibrations with slope of calibration and intercept were estimated using linear regression.

 
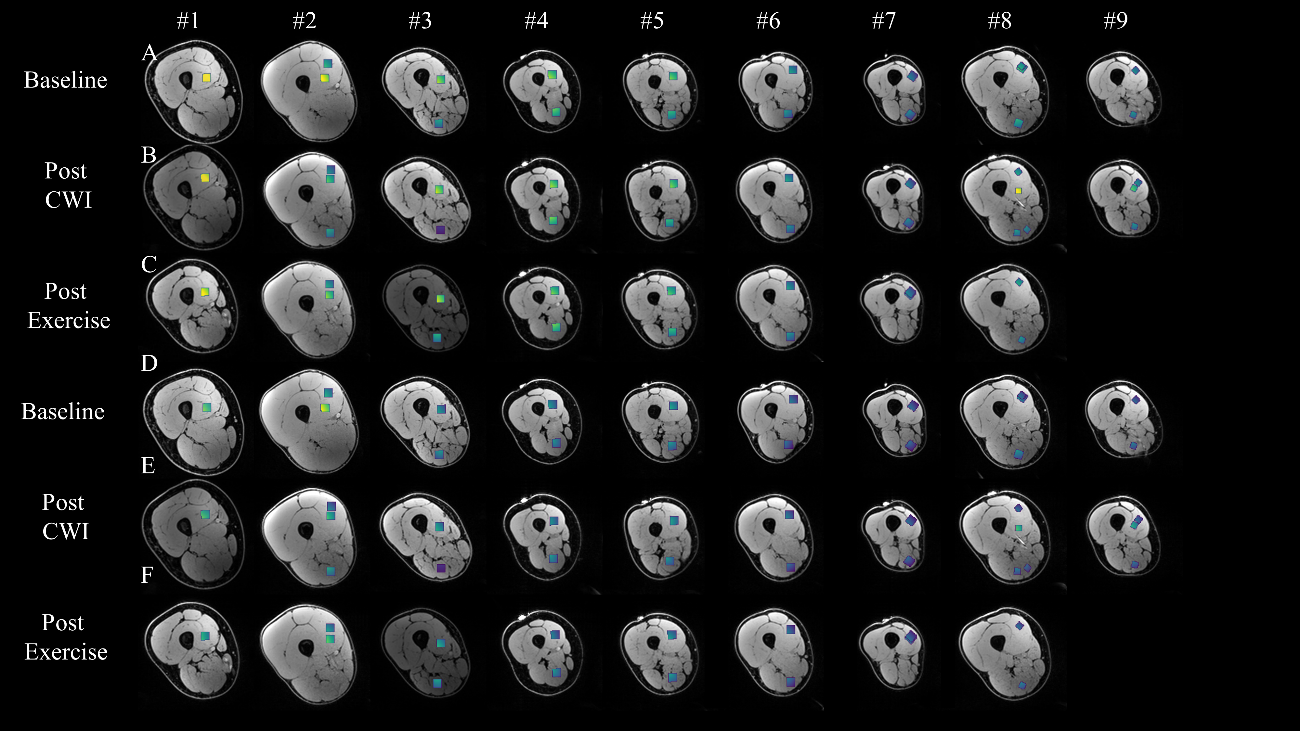

Supplementary figure 2:  VOI location (D) for the Baseline (A,D), post CWI (B,E) and post exercise (C,F) MRI exam overlaid on an axial T1 MRI image. The distance map within each voxel is indicated either computed from the skin surface (A,B,C) or from the muscle surface (D,E,F).

| **Volonteer** | **1** | **2** | **3** | **4** | **5** | **6** | **7** | **8** | **9** |
| --- | --- | --- | --- | --- | --- | --- | --- | --- | --- |
| **Age** | 27 | 26 | 61 | 30 | 39 | 26 | 49 | 26 | 45 |
| **Sexe** | M | M | M | F | M | M | F | M | M |
| **Size (m)** | 1.8 | 1.83 | 1.8 | 1.64 | 1.69 | 1.72 | 1.71 | 1.82 | 1.8 |
| **Weight (kg)** | 90.0 | 83.0 | 80.0 | 64.0 | 70.0 | 60.0 | 54.0 | 83.0 | 70.0 |
| **Volume total (cm³)** | 3926.4 | 4190.1 | 3211.1 | 3009.9 | 2883.8 | 2539.9 | 2124.9 | 3456.6 | 2327.5 |
| **Muscle volume (cm³)** | 2448.2 | 3673.9 | 2432.6 | 1586.4 | 1927.8 | 1826.9 | 1426.7 | 2517.0 | 1563.8 |
| **Fat volume (cm³)** | 1358.8 | 396.7 | 647.0 | 1340.6 | 859.2 | 619.3 | 582.9 | 823.4 | 651.1 |
| **Bone volume (cm³)** | 119.3 | 119.4 | 131.5 | 82.9 | 96.8 | 93.7 | 115.3 | 116.1 | 112.6 |
| **Ratio Muscle** | 0.624 | 0.877 | 0.758 | 0.527 | 0.668 | 0.719 | 0.671 | 0.728 | 0.672 |
| **Ratio Fat** | 0.346 | 0.095 | 0.202 | 0.445 | 0.298 | 0.244 | 0.274 | 0.238 | 0.28 |

Supplementary Table 1: Physical characteristics of the volunteers. Muscle, fat and bone volume are determined by segmenting the T1-weighted Dixon images. Fat fractions are also determined from the same sequence. Each volunteer's sport and level of training is reported.
